# Potential yield and food provisioning gains from rebuilding the world’s coral reef fish stocks

**DOI:** 10.1101/2025.05.06.652379

**Authors:** Jessica Zamborain-Mason, Joshua E. Cinner, Aaron MacNeil, Maria Beger, David Booth, Sebastian C.A. Ferse, Christopher Golden, Nicholas A. J. Graham, Andrew S. Hoey, David Moulliot, Sean R. Connolly

**Affiliations:** College of Science and Engineering, James Cook University, Townsville, Queensland, Australia; Department of Nutrition, Harvard TH Chan School of Public Health, Boston, MA, USA; Department of Environmental Health, Harvard TH Chan School of Public Health, Boston, MA, USA; Thriving Oceans Research Hub, School of Geosciences, University of Sydney, Camperdown, NSW, 2041 Australia; Department of Biology, Dalhousie University, Halifax, NS B3H 3J5, Canada; School of Biology, Faculty of Biological Sciences, University of Leeds, Leeds, LS2 9JT, United Kingdom; Centre for Biodiversity and Conservation Science, School of the Environment, The University of Queensland, Brisbane, QLD 4072, Australia; School of Life Sciences, University of Technology Sydney 2007 Australia; Leibniz Centre for Tropical Marine Research (ZMT), 28359 Bremen, Germany; Faculty of Biology & Chemistry (FB2), University of Bremen, 28359 Bremen, Germany; Faculty of Fisheries and Marine Sciences, IPB University, Bogor 16680, Indonesia; Lancaster Environment Centre, Lancaster University, Lancaster, LA1 4YQ, UK; MARBEC, University of Montpellier, CNRS, Ifremer, IRD, Montpellier, France; Smithsonian Tropical Research Institute, Panama City, Panama

**Keywords:** Food security, Multispecies fisheries, Coral reef fish, Sustainable yield, Recovery potential

## Abstract

Many coral reefs have fish stocks that are depleted below the level at which sustainable production is maximized. Lower production means that millions of people are losing out on potential food, income and livelihoods. Rebuilding these stocks to maximize sustainable production can contribute towards ending hunger and malnutrition but requires active and effective fisheries management. Yet, for fish stock recovery plans to be implemented, recovery benefits, targets and timeframes need to be quantified. Here, using 1211 individual reef sites and 23 jurisdictions identified globally as being below maximum sustainable production levels, we show that reefs have the potential to increase sustainable yields by nearly 50% if allowed to recover towards their maximum production levels. For individual jurisdictions this recovery represents from 20,000 up to 162 million additional sustainable servings of reef fish per year in comparison to current sustainable production, meeting recommended seafood intake for up to 1. 4 million additional people a year. However, such growth and food provisioning will require fish stocks to double their standing biomass (increase by a median of 32 t/km^2^). Recovery timeframes range from 6.4 years under the most stringent scenario (a seascape moratorium) to 49.7 years under the maximum harvest scenario that results in recovery. We find that locations with the greatest potential for sustainable gains in yield are among those with the greatest food and micronutrient deficiencies, underscoring both the challenges and opportunities in recovering fish assemblages to achieve their maximum sustainable potential.

**Significance Statement:** Coral reef fisheries are a critical food source for people throughout the tropics. However, most reefs around the globe have fish biomass values below those enabling maximal sustainable production, risking food availability, income, and livelihoods. Rebuilding fish assemblages can increase sustainable food supplies, and, if these are well distributed, directly contribute towards enhanced food security. We show that recovering fish stocks on coral reefs can significantly increase the number of sustainable fish servings produced per year and the number of people meeting fish intake recommendations, particularly for countries with high malnutrition. Our study highlights the sustainable food provisioning potential of recovering reef fisheries and quantifies how much recovery would be needed and the time such recovery would take.

## Introduction

Multispecies coral reef fisheries often operate in locations with compromised food security, employment, and income (1-3). In such locations, reef resources can be a readily accessible (4) source of food and nutrients, contributing to nutritional intake both directly, as key sources of vitamin B_12_, haem iron, retinol, niacin, and protein (5) and indirectly by generating income that allows people to purchase other nutrient-dense foods (6). Thus, managing reef resources to maximize sustainable food provisioning is a path towards meeting key sustainable development goals such as “Zero Hunger” (7). However, recent global assessments of coral reef fisheries have shown that most reefs have fish biomass values below those that would maximize sustainable production and are not delivering their full yield potential (8, 9). For example, 64% of 1903 exploited reef sites and 47% of 49 tropical jurisdictions assessed globally had reef fish stocks that were below 50% of their estimated unfished biomass (9), a common reference point at which yields are expected to be maximized (10). Having assemblages below reference point values increases the risk of stock collapse (10), decreases economic returns (11), imperils ecosystem functioning (8, 12, 13), and can jeopardise fishery yields and food security (14-16). Low fish biomass levels means that the millions of people that depend on reef ecosystems (17) are losing out on potential sustainable food supplies, income, and livelihood opportunities.

If fished populations become depleted below reference point levels aimed at maximizing sustainable production, the main goal of fisheries management is often to recover populations back to those levels (10, 18). For multispecies fisheries, key reference points can include those that produce “multispecies maximum sustainable yields” (MMSY (19)) or “pretty good multispecies yields” (PGMY; defined as a catch that is ≥0.8 of MMSY (20, 21)). However, in many locations, such as those where multispecies coral reef fisheries operate, managing fish stocks to achieve recovery remains a major challenge (18). Recovery costs (22), together with socio-economic and management constraints (1, 23), and a lack of defined context-specific recovery targets and timeframes (8), tend to outweigh the unknown or uncertain benefits from recovery, hindering the design of, and compliance with, effective fisheries management plans (24, 25).

Quantifying recovery targets and timeframes is necessary to effectively implement recovery management plans (10, 26). Additionally, quantifying foregone sustainable yields presents a means to evaluate costs of recovery plans against potential yield and food provisioning gains from recovered stocks (27). Yet, the extent to which rebuilding fish stocks on coral reefs can contribute towards ending hunger and malnutrition is currently unknown. Here, using data from 1211 coral reef sites and 23 jurisdictions identified globally as having fish assemblages below 50% of their unfished biomass (Methods (9)), we: (i) quantify the sustainable yield lost due to having fish stocks below levels aimed at maximizing sustainable production (Fig. 1); (ii) highlight the benefits of recovering fish assemblages in terms of food provisioning; (iii) estimate the standing biomass increases required to recover assemblages to MMSY (i.e., B_MMSY_) and biomass levels that produce PGMY (i.e., B_MMSY_ and B_PGMY_); and (iv) estimate the time it would take to recover multispecies fish assemblages to those levels.

**Figure 1.**
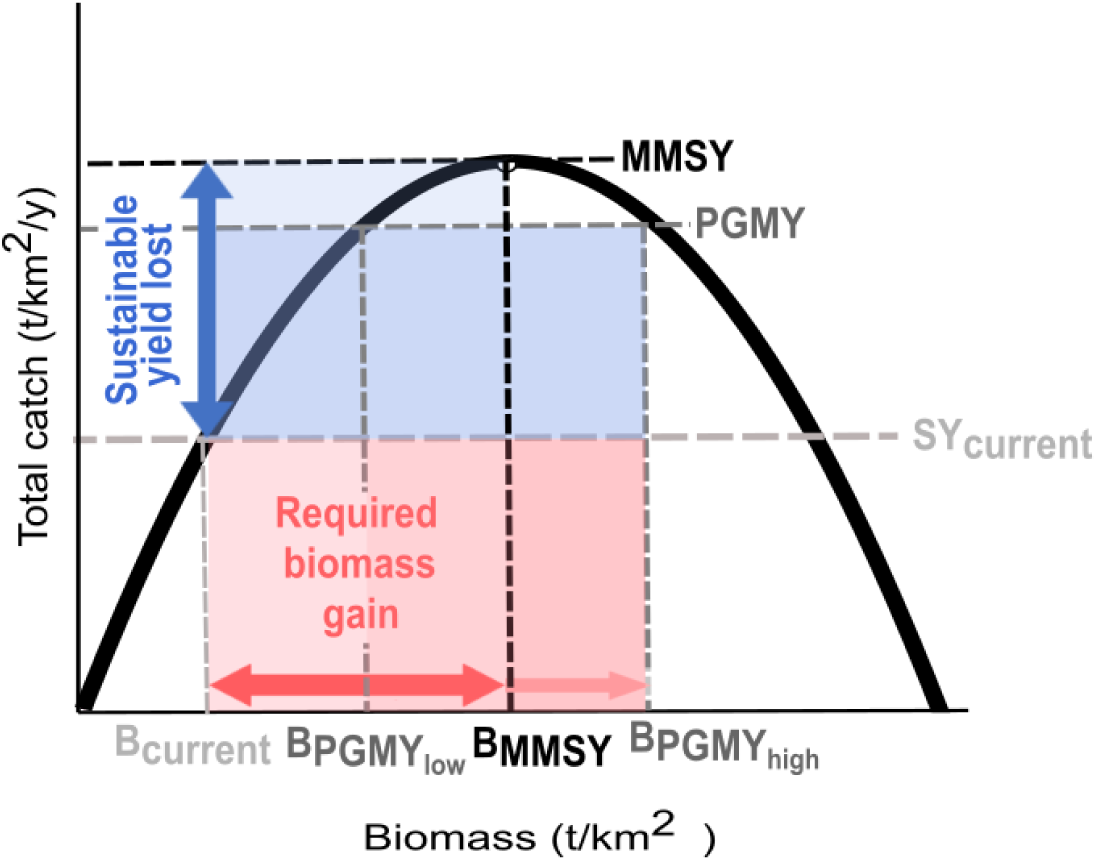
Theoretical relationship between sustainable yield losses and required biomass gains in coral reef fisheries. Conceptual surplus production curve diagram exemplifying how to quantify (i) the sustainable yield lost from having fish assemblages below biomass levels aimed at maximizing sustainable production B_MMSY_ (B_current_<B_MMSY_) and (ii) the required biomass increases to recover assemblages to either maximum production values (MMSY) or pretty good multispecies yields (PGMY). The x axis represents the standing fish assemblage biomass and the y axis the sustainable total catch or estimated surplus production for the whole multispecies assemblage. The solid line represents the expected surplus production along a gradient of biomass (assuming a Graham-Schaefer surplus curve for the multispecies assemblage, according to which MMSY is achieved at biomass levels corresponding to 50% of unfished biomass). SY_current_ is the expected long-term yield (i.e., surplus) from a hypothetical multispecies assemblage given its standing biomass (B_current_); B_MMSY_ is the estimated biomass at which MMSY is achieved; and B_PGMY_ are the estimated biomass at which PGMY are attained (below and above B_MMSY_: B_PGMY,low_ and B_PGMY,high_, respectively). Note that we are not comparing current yields (i.e., how much is being caught on a reef) with potential maximum sustainable yields (i.e., how much could be caught sustainably if managing assemblages to maximize production) but instead how much sustainable yields could increase from their current hypothetical sustainable levels,

## Results and Discussion

### Sustainable yield lost from stocks below B_MMSY_

We quantified the sustainable yield lost from depleted coral reef fish assemblages for each site and jurisdiction classified below B_MMSY_ as the difference between the estimated MMSY and the estimated surplus production from the standing fish biomass (Fig. 1)– based on location-specific sustainable reference points estimated from a multispecies Graham-Schaefer surplus production model (Methods; (9)). We found that, globally and based on median values, reefs classified as below B_MMSY_ are provisioning 68% of their maximum (Fig. S1; (28)) and have a sustainable production deficit of 1.9 t/km^2^/y, but with substantial among site variation (from 0.1 to 18.8 t/km^2^/y; site-specific posterior medians; Fig. 2A-C). This deficit represents a potential increase of ∼47 [1.7 – 14574.4] % (median [min-max] across sites) in long-term yields compared to current surplus levels if stocks were allowed to recover to maximum production (B_MMSY_). Converting these yields to 100g fish servings and to the number of people potentially meeting fish intake recommendations for cardiovascular health (Methods; (29)), we show that an additional 9088 [276 – 93255] reef fish servings could be produced yearly per square kilometre of coral reef, or 77 [2 – 791] additional people per square kilometre of reef could meet their yearly fish intake recommendations. A smaller increase of 0.7 [-1.8 – 14.7] t/km^2^/y or 17 [-18.4 – 11790.7] % would be possible if stocks were allowed to recover to PGMY values (e.g., B_PGMY,low_), which translates into 3361 [-9160 – 69446] fish servings/km^2^/y or 29 [-78 – 621] people meeting yearly fish intake recommendations (with negative values indicating such sites are already between B_PGMY,low_ and B_MMSY_, and thus producing > PGMY).

**Figure 2.**
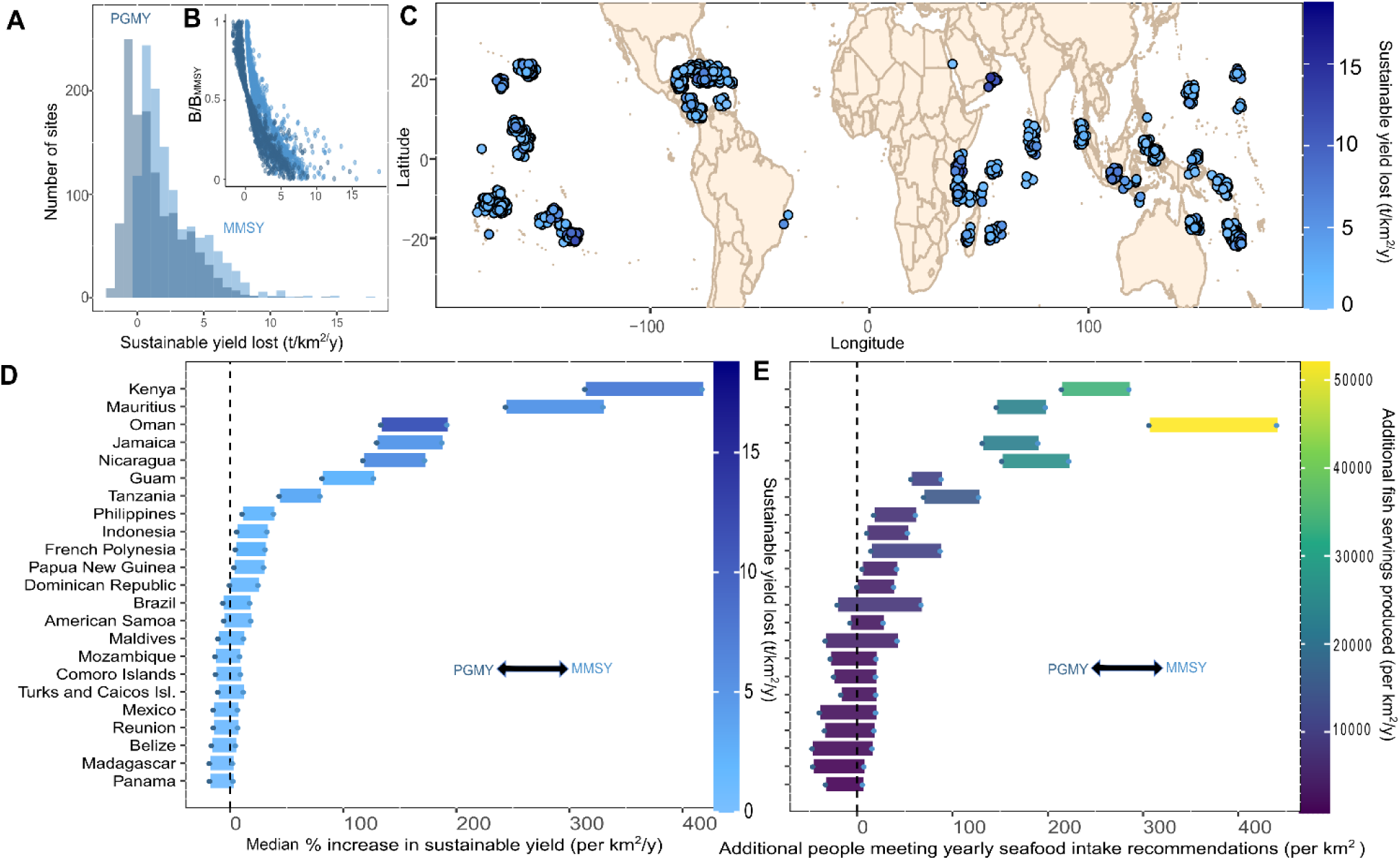
Per unit area sustainable yield lost from coral reef assemblages. (**A**) Posterior median sustainable yield loss for coral reef sites open to fishing classified as being below B_MMSY_. Colours represent losses based on MMSY (light blue) or PGMY (dark blue). Negative values in the PGMY distribution means those sites are already within PGMY values (i.e., between B_PGMY,low_ and B_MMSY_). (**B**) Relationship between expected sustainable yield lost and site-specific biomass status (B/B_MMSY_). Points are posterior medians for each exploited site classified as below B_MMSY_. Different colours are the same sites but for MMSY and PGMY. (**C**) Map of sampled fished sites below B_MMSY_, colour-coded by the median sustainable yield lost based on MMSY. Points are jittered to add clarity. (**D**) Median % increase in sustainable yields expected per jurisdiction if recovered to MMSY (upper limit of horizontal bar) or PGMY (lower limit of horizontal bar) in comparison to current estimated sustainable production. Only jurisdictions which have median estimated biomass values below B_MMSY_ are included (biomass values are weighted optimistically assuming the proportion of marine protected areas in a jurisdiction are at unfished biomass values; Methods). Bars are coloured by the median sustainable yield lost based on MMSY. (**E**) Additional people potentially meeting yearly seafood intake recommendations per square kilometre for each jurisdiction based on median values. Bars are coloured by the potential additional number of fish servings produced per km_2_ if reef assemblages are recovered to MMSY. Note that in **D-E** if bars overlap the horizontal dashed line (∼0 % increase) it means that jurisdictions median biomass for exploited reefs is estimated to be within B_PGMY,low_ and B_MMSY_ (i.e., they are already estimated to be producing > 80% of MMSY, so meeting 80% can result in negative numbers).

Lost sustainable yields and food supplies are widespread among all geographic basins (Fig. 2C). However, the greatest estimated yield losses occur in locations where fish assemblages are most depleted (Fig. 2B), meaning they have the greatest potential long-term gains in yield. For example, at a jurisdiction scale, Kenya, Mauritius or Oman – which have median biomass values below 10% of their estimated unfished biomass (9) –could increase their median per-unit-area sustainable catches by 418, 330, and 193 %, respectively, if stocks were allowed to recover to levels that produce MMSY, with slightly lower, but still substantial, increases with recovery to PGMY (314, 244 and 134 %; Fig. 2D). For these specific jurisdictions, increased yields to MMSY could translate into >23000 additional sustainable reef fish servings (each 100g) produced per year per square kilometre of reef or > 198 additional people meeting yearly fish intake recommendations per square kilometre of coral reef in comparison to current sustainable production (Fig. 2E).

### Food and potential nutrition benefits of recovering stocks on coral reefs

Overall, by scaling sustainable yield gains and fish servings to their corresponding reef area open to fishing (Methods), we found that jurisdictions are missing out between 3.9 to 32389.6 t/y (jurisdiction-specific medians) by having fish assemblages below MMSY values, and thus have the potential to produce from 0.02 to 161.95 million additional fish servings a year compared to their current estimated production (Fig. S2). These numbers mean that, dependent on the jurisdiction, from 166 to 1373208 additional people could meet yearly fish intake recommendations from coral reef fish recovery (Fig. 3A). For nations such as French Polynesia people meeting fish intake recommendations from gained yields represents 97% of the coastal population, and for others such as Tanzania, Maldives and Mauritius, > 20% of the coastal population could meet yearly fish intake recommendations from the gained yields (Fig. 3A; Table S1).

**Figure 3.**
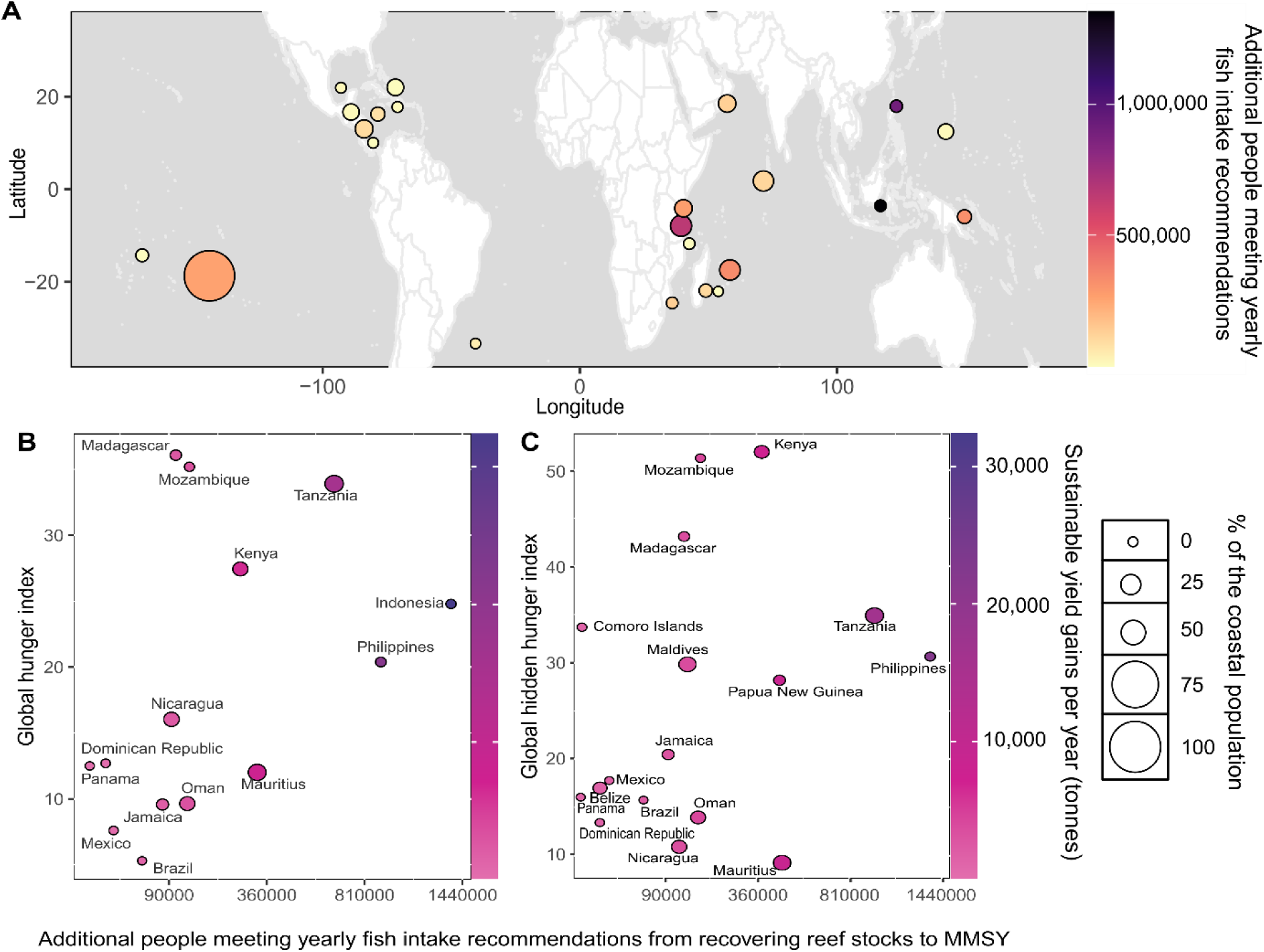
Tackling hunger and micronutrient deficiencies by rebuilding reef fish stocks to maximum production levels. (**A**) Jurisdiction-specific estimated number of additional people that could meet yearly fish intake recommendations from the sustainable yields generated from recovering fish assemblages to MMSY values. (**B-C**) Relationship between the estimated number of people meeting yearly fish intake recommendations from recovering reef fish assemblages to MMSY values and (**B**) global hunger and (**C**) hidden hunger indices. Each point is a jurisdiction classified as below B_MMSY_ with available index information (i.e., jurisdictions in panel **A** that are not highlighted in panels **B-C** did not have these indices available, Methods). Points are colour-coded by the estimated yield gains based on jurisdiction-specific per-unit-area yield gains to MMSY extrapolated to the estimated reef area open to fishing (excluding the percentage of area estimated to be protected; Methods). Point size in **A-C** is scaled based on the % of coastal population that could meet yearly fish intake recommendations from gained yields (i.e., additional people who would benefit from recovering fisheries as a percentage of the coastal population). See Table S1 for actual numbers.

Based on the current status of fish stocks, jurisdictions can provide from 0.26 (i.e., Reunion) to 483.58 (i.e., Indonesia) million sustainable reef fish servings per year, meeting the yearly fish intake recommendations for up to 4.1 million people (i.e., Indonesia; Table S1). For jurisdictions such as French Polynesia, their estimated sustainable production already meets the seafood intake recommendations for > 100% of the coastal population. However, for others (i.e., Jamaica, Philippines, Dominican Republic, Indonesia, Panama, Reunion, Mexico, Brazil and Kenya) current estimated sustainable production covers much lower percentages of the costal population (<5%; Table S1). Allowing fish stocks to recover to MMSY can significantly benefit jurisdictions. Indonesia, for example, could provide enough yearly seafood intake recommendations for a total of ∼5.5 million people if their reef fish stocks were managed at MMSY. Similarly, >50% of the coastal population from Tanzania, Turks and Caicos, Madagascar, Belize and Maldives could meet yearly intake recommendations from recovered stocks. Recovering fish stocks to MMSY could allow some jurisdictions to double (i.e., Guam), triple (i.e., Jamaica, Nicaragua and Oman) or increase above four (i.e., Mauritius) and five (i.e., Kenya) fold the percent of coastal population meeting yearly fish intake recommendations (Table S1). Increased sustainable yields from recovering reef fish stocks, thus, can substantially contribute to enhanced food security.

By matching the number of additional people meeting intake recommendations to jurisdiction-specific hunger and hidden-hunger indexes (Methods), we found that places with higher human hunger (i.e., hunger and undernourishment (30, 31)) and hidden hunger (i.e., micronutrient deficiencies (32, 33)) are within jurisdictions with the greatest potential sustainable yield and food supply gains (Fig. 3; Spearman’s correlation of ∼ 0.5 and 0.3, respectively). For example, jurisdictions with “Alarming” and “Severe” hunger values (i.e., Madagascar, Mozambique, Tanzania, Kenya, Indonesia and Philippines), which have inadequate food supplies, high child undernutrition and mortality (30, 31), are missing out on 2545 to 32390 t/y of sustainable reef fish production by having reef stocks below maximum sustainable production values, equivalent to 12.7 to 161.9 million additional 100g reef fish servings per year or 107914 to 1373208 additional people potentially meeting yearly fish intake recommendations (Fig. 3B).

Similar conclusions apply to jurisdictions with “Alarming” and “Severe” micronutrient deficiencies in pre-school children (i.e., Madagascar, Mozambique, Tanzania and Kenya; (32)), which are losing 2545 to 15814 tonnes of reef fish, 12.7 to 79.1 million reef fish servings per year, and failing to meet the fish intake recommendations for 107914 to 670473 number of people (Fig. 3C). Management of wild fisheries to maximum sustainable production values is one of the main pathways to increase food supplies from the sea (16) and a direct way to tackle global micronutrient deficiencies (3, 14). Tropical marine fish are rich in calcium, iron, vitamin A and zinc in comparison to other marine fish species (3) and reef fish, in particular, can be the main source of vitamin B_12_, iron, niacin and vitamin A in the diet of tropical coastal communities (5). Thus, by rebuilding reef assemblages and retaining the increased catches within the jurisdiction, one can increase sustainable food quantities and, if distributed equitably, partially or fully address nutrient deficiencies using local fish stocks. This highlights the potential of fisheries recovery plans to contribute towards policies that aim to tackle pervasive food insecurity in tropical developing regions.

### Recovery targets and timeframes

Sustainable yield gains and their benefits can only be realized if depleted fish stocks are allowed to recover, which in turn requires temporary reductions in catch to below current production levels, with larger reductions typically facilitating faster recovery. Consequently, we estimated the site-specific and jurisdiction-level standing biomass increases that would be required to obtain (i) the biomass at which MMSY is expected under a Graham-Schaefer surplus production model (i.e., B_MMSY_; Fig. 1), and (ii) corresponding PGMY biomass levels (i.e., B_PGMY,low_ and B_PGMY,high_). We found that, to reach maximum production levels at B_MMSY_, reef sites would have to double their standing biomass by a median of 32 t/km^2^, but with substantial among-site variation depending on the state of depletion [from 0.4 to192.9 t/km^2^]. Lower biomass increases (6 [-45.2 – 102.2] t/km^2^) would be required to reach PGMY levels below B_MMSY_ and larger biomass increases (62 [17.2 – 283.7] t/km^2^) would be required to reach PGMY levels above B_MMSY_– a target that entails less risk of imposing catches that overshoot true maximum sustainable yields (20) and is associated with increased biodiversity and ecosystem benefits (e.g., 8, 9). Jurisdictions like Oman, Kenya or Mauritius, for example, are among those with less current surplus production and major potential benefits per unit area but also require substantial standing biomass recovery to achieve them: 132.9, 78.4 and 55.4 t/km^2^ to reach B_MMSY_ and ∼ 0.5 or 1.5 times that to reach B_PGMY,low_ and B_PGMY,high_, respectively (Fig. 4C).

**Figure 4.**
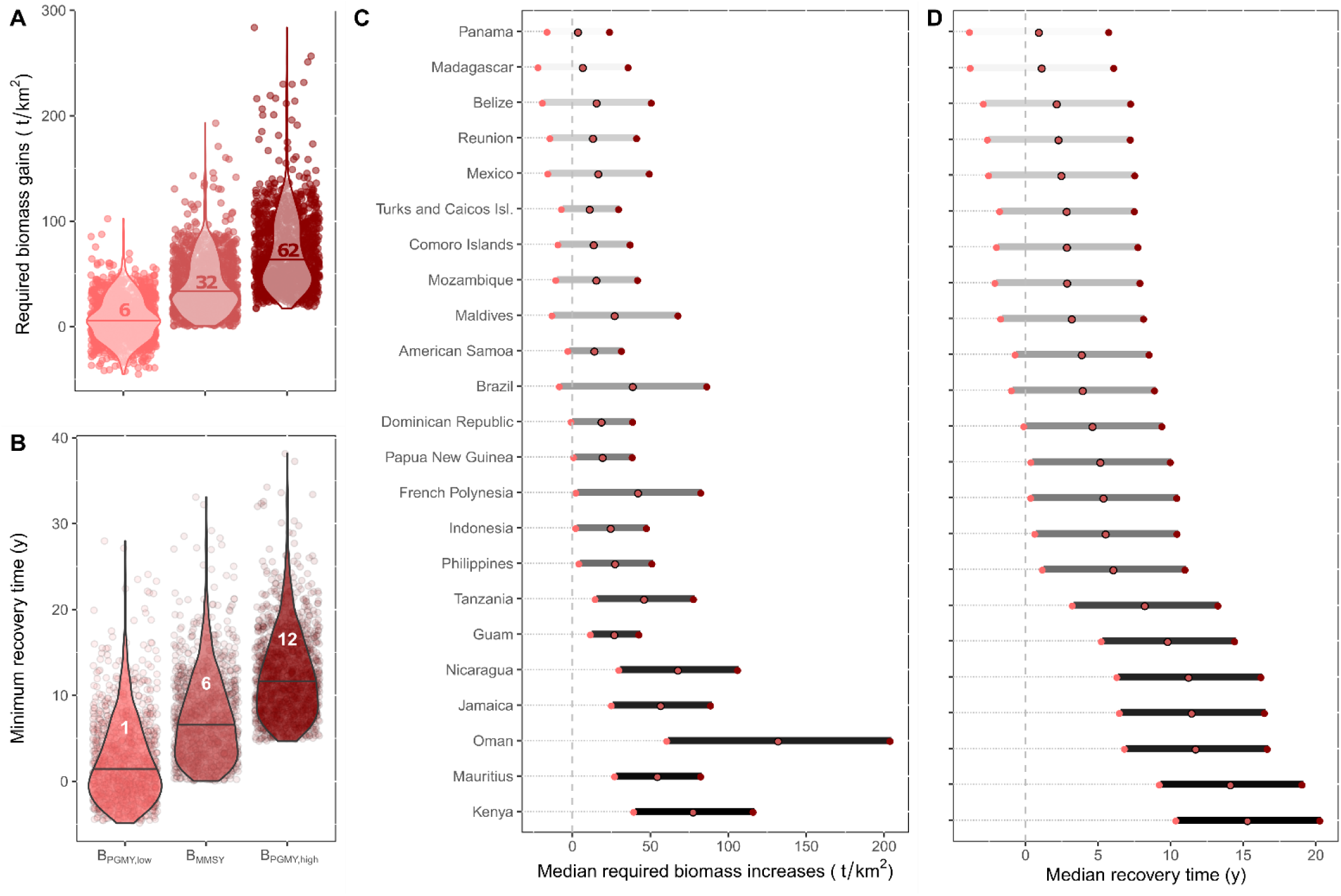
Median required biomass recovery and timeframes. (**A**) Distributions of posterior median site-specific biomass increases required to achieve site-specific B_PGMY,low_, B_MMSY_ and B_PGMY,high_ levels, respectively. (**B**) Distributions of site-specific recovery timeframes (posterior medians) under the most stringent seascape-wide moratorium scenario (i.e., minimum recovery time) to recover stocks below B_MMSY_ to their estimated site-specific B_PGMY,low_, B_MMSY_ and B_PGMY,high_ levels, respectively. Numbers in (**A-B**) are median values for all sites. (**C**) Median required biomass gains per jurisdiction. (**D**) Estimated jurisdiction-specific recovery timeframes under the most stringent seascape-wide moratorium scenario (i.e., minimum recovery times). Horizontal bars in (**C-D**) represent the biomass gains or recovery times required to reach B_PGMY,low_ (left limit) and B_PGMY,high_ (right limit) given a jurisdiction’s current sustainable surplus and standing stock biomass, whereas the middle point represents B_MMSY_ conditions. Colours in (**C-D**) are based on the biomass status (B/B_MMSY_) with darker meaning worse (more below B_MMSY_). See Fig. S3 for recovery timeframes with harvesting.

The key impediment to rebuilding stocks towards target biomass values lies in the near to medium-term costs (e.g., loss of catch) to be borne by resource users. We estimated that, when fishing for recovery, reef sites can take from 6.4 to 49.7 years to reach maximum production biomass levels (Fig. S3), More specifically, in the most stringent and fastest recovery scenario (10), where seascape-wide fishing moratoria are in place on otherwise undisturbed reefs, locations estimated to be below B_MMSY_ would take a median of 6.4 years [0.03 – 33.1; dependent on site] to recover towards assemblage-level maximum production (B_MMSY_), 1.3 [-4.9 – 27.9] years to reach B^PGMY,low^ values and 11.6 [4.7-38.2] years to reach the PGMY targets above B_MMSY_ (Fig. 4B). However, the estimated recovery timeframes increase as a larger proportion of the standing stock is harvested (e.g., Fig. S3), with recovery times reaching up to 49.7 [3.7-106.7] years under the maximum harvesting regime that allows for recovery to B_MMSY_ (Fig. S3). For heavily depleted jurisdictions like Oman, Mauritius or Kenya, we estimate that recovering stocks to B_MMSY_ can take up to 70 years under the maximum harvesting regime that allows for recovery (Fig. S3), with a minimum of 11 years in a scenario where the entire seascape is closed to fishing (Fig. 4D)-an approach that contrasts with the current practise of single reserve patches within fished seascapes that likely export biomass (e.g., 8, 34). Such seascape-scale moratoria are not socially desirable, feasible or economically viable (11) for most reef fishery locations, especially where dependence on fisheries for food and nutrition security is high. Consequently, our seascape fishing moratoria are not intended as a policy goal, but rather an estimation of minimum recovery timeframes to help inform alternative rebuilding actions.

### Challenges and opportunities of realizing recovery

In contrast to many well-studied fish stocks where fisheries management strategies have been implemented to exploit stocks at sustainable levels (18, 19, 35), most coral reef multispecies fisheries lack information about the ecological, social, and economic costs and benefits of different recovery pathways and their trade-offs. Management measures such as gear and effort restrictions (8), spatial or temporal closures (36), size and species bans (10), the displacement of effort to other available resources (e.g., small pelagic resources or mariculture (37)), or a combination of tools (e.g., 11) have the potential to allow reef assemblages to recover and realize their sustainable yield and food supply gains, yet the most-effective pathway will be context-specific to metapopulation structure, species composition and other features of the socio-ecological system.

Ecologically, the lack of spatio-temporal data on recovery limits the extent to which we understand the functional form of the surplus production curve for whole assemblage biomass in different locations (9) and how this is impacted by environmental change, species composition (e.g., trophic turnover (38, 39)) or novel reef assemblages (40). For example, here we estimated recovery timeframes for the multispecies assemblage using community growth rates inferred from different reserve sites of different ages (i.e., a space-for-time substitution (9)) and assuming production peaks at 50% of unfished biomass values of a symmetric production curve. However, timeframes and biomass recovery targets to achieve maximum production could differ if the fish assemblage recovers at a different rate or the assemblage production function peaks at different biomass values.

Recovery and its timelines can be influenced by a range of factors, including trophodynamics (39), habitat impacts (26, 36), the state of depletion (35), and changing environmental contexts (26, 35, 41). For coral reef fish, in particular, how biomass recovers is not solely correlated with metrics related to fishing intensity (42), suggesting that cumulative impacts on the seascape will likely affect recovery (43). Furthermore, biomass on reefs is predicted to decrease with increased projected thermal stress (44, 45), but given that most fisheries are not effectively managed to maximize production, how such expected decreases will impact recovery to maximum production targets, or the targets themselves, is not straightforward (46). Rebuilding assemblages to levels that correspond to 50% of unfished biomass (e.g., B_MMSY_) or higher (e.g., B_PGMY,high_) is associated with stock and ecosystem benefits such as increased mean fish length and species richness, and enhanced ecosystem functioning (8,9,12), which are likely to feedback as long-term socio-economic gains (47). Yet, a better understanding of recovery in terms of how community growth rates vary spatially and temporally (41) or the species available for harvest at different recovery phases (38, 39) will help fine-tune these benefits and allow to dynamically adjust targets and timelines under different contexts (48).

Food provisioning benefits will only contribute to food security if sustainable reef fish production is accessible to those who need it most. For example, benefits of recovery may be jeopardized if locations have social, economic, or institutional barriers preventing equitable access to them (4,49-51). While stock recovery is the first step towards enhanced sustainable food supplies, pausing or reducing fishing to promote recovery will impact people’s reef fish supplies and will require alternate food sources from agriculture, mariculture or offshore resources (14) that ensure people have access to adequate amounts of nutritious foods. Reef fish are an important source of food and nutrients for people that depend on reef-based food systems (5), so promoting a reduction in reef fish consumption to enhance recovery will require active food-system interventions that are able to overcome established cultural and supply links to reef fish. This is especially the case for locations currently overfishing their reef fish stocks beyond sustainable levels (9), where current yields being harvested surpass the estimated surplus production or MMSY, and where maximum sustainable production actually results in a decrease in the yearly number of humans meeting fish intake recommendations in comparison to unsustainable yields (e.g., Indonesia, Philippines, Mexico, Dominican Republic, Jamaica, Nicaragua, Oman and Panama; Fig. S4). Locations with large and fertile arable lands and diverse food sources might be compatible with temporary or well-placed sea closures, but many small island states without arable land may require ocean (16) or trade-based solutions besides gradual decreases in fishing effort to realize their potential reef-fish benefits.

Recovering coral reef fish populations to maximum production biomass also requires considerations of multiple time horizons that impact projected returns on investment (2). For instance, successfully realizing recovery requires active fisheries management, monitoring and enforcement (10). This will generate economic gains in the long-term (52), however establishing fisheries reforms and monitoring their effectiveness will also require short to medium-term economic investments, and evaluating the trade-offs of implementing different management measures (53). Reductions in fishing, even if temporally, will require alternative innovative livelihood options such as tourism, agriculture, or mariculture (37, 54) and institutional systems that provide the right incentives (e.g., economic) to displaced resource users (55). Their success, will be context-specific. For example, locations with high tourism potential might be compatible with fishing bans where tourism is not reliant on reef fisheries (56) and tourism benefits are distributed among those affected by recovery plans. For places with low tourism potential a transition to sustainable agriculture or mariculture might be more advantageous if conditions and infrastructure permit it (54). Economic incentives for individual operators to recover their fish assemblages might be possible for some high development countries. However, given the lack of strong institutions in most locations where reef fisheries operate (1), many reef locations may require international support systems that support recovery and their complex socio-ecological landscape (e.g., 57).

Another social dimension to consider is employment and the social cost of employment displacement. For locations that have high percentage of fishers within the population, recovery might translate into excessive employment costs, whereas for others employment displacements might be relatively minimal and feasible with incentives. However, reefs and their resources are important to communities in a number of ways beyond food, income, and employment (58), supporting deep cultural values, identity, and wellbeing (59-61). Having healthy resources is likely to benefit fishers multiple ways, but cessation of fishing to promote recovery will bring context-specific costs that will have to be considered at relevant scales to successfully achieve recovery (62).

## Conclusions

Two key, interdependent United Nations Sustainable Development Goals are to “conserve and sustainably use marine resources for sustainable development”, as well as to “end hunger” (https://sustainabledevelopment.un.org; 63). Here, we have quantified the sustainable fish yields lost as a consequence of reef fish assemblages being below maximum production levels and the potential sustainable long-term food provisioning gains that can be achieved from their recovery. Yet to realize those potential benefits, reef fish assemblages need to be substantially recovered. Locations that most suffer from hunger and micronutrient deficiencies are within those with the greatest potential for sustainable yield gains, yet they are also where assemblages are most depleted, where more and longer recovery will be required, and thus where it is probably hardest to implement fisheries management plans to promote recovery. We provide benchmark recovery targets and timeframes against which to judge alternative recovery scenarios. The challenge now is finding effective pathways to recovery for each context even under climate change, ensuring people have access to nutritious foods. This will likely require increased local knowledge and participation, creation of alternative livelihood and food supplies, long-term planning, continuous monitoring, and financial and other support mechanisms that allow tropical communities to recover their lost potential yields.

## Materials and Methods

### Yield lost

Site-specific and jurisdiction-scale standardized reef fish biomass estimates and sustainable production curves were obtained for coral reef fish assemblages using the same approach as a recent global assessment that calculated site-specific multispecies sustainable reference points for coral reef fish (Table S2) based on environmental conditions (9). That study used a total of 2053 reef sites worldwide: 150 were high-compliance marine reserves of different ages or remote uninhabited reefs that were critical to infer the unfished biomass for reef fish assemblages, how the unfished biomass varies with environmental context (i.e., coral cover, ocean productivity, sea surface temperature, and whether the reef is an atoll), and how reef fish populations grow in the absence of fishing toward unfished biomass (i.e., community biomass growth rate); the remaining 1903 reefs informed sampling parameters (e.g., related to census method, sampling area, habitat type, and depth of survey), and their biomass was used to assess the status of exploited fish assemblages at a site and/or jurisdiction-level scales (e.g., contrasting the biomass per unit area and/or annual catch per unit area to the estimated reference points aimed at maximizing sustainable production (9)). The study used different surplus production curves, all of which fit the recovery data relatively well. Here we used global estimates based on an aggregate Graham–Schaefer surplus growth model in which the whole assemblage is treated as a single population (64, 65), and a symmetric (logistic) population growth curve whose productivity (and thus sustainable yields) peaks at 50% of unfished biomass is assumed (66, 67).

As the aim of this study was to quantify the expected sustainable yield lost from exploited assemblages below levels aimed at maximizing sustainable production, from the 1903 reef exploited reef sites assessed (i.e., defined as openly fished and restricted), we only included 1211 (64%) sites that, based on median estimates, had reef fish biomass values below 50% of their estimated unfished biomass (i.e., those where biomass was estimated to be below the biomass at which multispecies maximum sustainable yields or B_MMSY_ is expected to be achieved). Similarly, from all jurisdictions with biomass data available (N=49), we only included jurisdictions that had median biomass values (weighted by the reported percentage of territorial waters protected) also considered to be below B_MMSY_ (N=23). These weighted biomass values per jurisdiction optimistically assume that the percentage of territorial waters that are protected from fishing are at unfished biomass values and exploited individual reef sites sampled within a jurisdiction are representative of jurisdiction-level conditions (9). For each site, we extracted (i) the estimated posterior marginalized biomass (estimated standing stock biomass standardized for methodological effects), and (ii) the estimated surplus production curve given a site’s environmental conditions (i.e., B_MMSY_, MMSY and the estimated surplus along a gradient of biomass). Note that for sites, we assume sampled reefs are representative of the conditions at the scale of metapopulation closure. Site-specific yield losses were calculated for each site as the difference between the estimated site-specific MMSY and the site-specific estimated surplus (e.g., Fig. 1). We also calculated proportions (surplus/MMSY), equivalent to the food provision index in ref (28) (Fig. S1).

Similarly, to estimate the median yield lost per jurisdiction (e.g., Fig 2C), we subtracted the median jurisdiction-level estimated surplus (based on median weighted biomass values and jurisdiction-specific surplus production curves) from the median jurisdiction-level MMSY. To estimate jurisdiction-level yearly potential yield gains, jurisdiction per-unit-area sustainable yield losses (i.e., potential gains) were extrapolated to the reef area open to fishing (68, 69), assuming the proportion of territorial waters protected within a jurisdiction (70, 71) was equal to the proportion of reef area within a jurisdiction closed to fishing.

Note that this is not comparing current yields (i.e., how much is harvested) to potential maximum sustainable yields (i.e., the maximum that can be harvested sustainably based on surplus production models), but instead current estimated sustainable yields (i.e., how much can be harvested sustainably based on the biomass of a site or jurisdiction and surplus production models) to potential maximum sustainable yields. A jurisdiction or site can be subject to overfishing, and in such a case, actual yields can surpass sustainable yields and even maximum sustainable yields. See Fig. S4 for a comparison to actual yields estimated over coral reefs (9) based on reconstructed reef fish landings.

### Fish servings and fish intake recommendations

Yearly potential yield gains of reef fish at the site and jurisdiction level were converted to 100 g fish servings by converting live weight estimated production to edible weights, conservatively assuming 50% of reef fish live weight produced (in metric tonnes) are edible (45; 72). We also calculated the number of people meeting yearly recommended seafood intake from estimated yield gains by combining edible weights with per capita weekly seafood intake recommendations of 226.796 grams (i.e., 8 oz (29)) and assuming 52 weeks in a year. These weekly intake seafood recommendations are based on long-chain omega 3 fatty acids concentrations in seafood and recommended intake to minimize the risk of cardiovascular disease (29). At a jurisdiction scale, additional servings and people meeting recommended intake values per year were estimated using jurisdiction-level yearly potential yield gains that account for jurisdiction-specific coral reef areas potentially open to fishing. Additional people meeting recommended fish intakes were then converted to percents of the coastal population using the estimated coastal population size within 30km of the coastline (calculated from Grided Population of the World for 2019; (73, 74)).

### Global hunger and hidden hunger indexes

Available jurisdiction-level global hunger and global hidden hunger in pre-school children indices were obtained from Our World in Data (https://ourworldindata.org/; (30-33)). The global hunger index was obtained for 2010 (consistent with most reef fish assemblage data (9)) and combined four indicators: Undernourishment, child wasting, child stunting and child mortality (30). The global hidden hunger in pre-school (age <5 years) index was obtained for the period 1999-2009 and is calculated as the average of three deficiency prevalence estimates weighted equally: preschool children affected by stunting, anemia due to iron deficiency, and vitamin-A deficiency (32). Note that these indices were not calculated for some jurisdictions in the selected time periods so these were excluded for this part of the analyses (e.g., French Polynesia). For the hidden hunger index, scores between 0 and 19.9 are considered mild, 20-34.9 moderate, 35-44.9 severe, and 45-100 are considered alarmingly high (33). For the hunger index, scores between 0 and 10 are considered low, 10-20 moderate, 20-35 serious, 35-50 alarming, and >50 extremely alarming (31). To categorize indexes consistently, we re-named “alarmingly high” to alarming, “Serious” to severe, and “Low” to mild in the hidden-hunger index.

### Required biomass gains and recovery timeframes

Location-specific required biomass gains to achieve B_MMSY_ or B_PGMY_ were calculated for each location (i.e., site or jurisdiction) as the difference between the estimated biomass target (B_PGMY,low_, B_MMSY_ or B_PGMY,high_) and the estimated biomass (e.g., Fig. 1). Although B_PGMY,low_ should not be considered a target (because it is below B_MMSY_ and imposes a higher risk of driving stocks to collapse), it may be useful from a food security perspective (e.g., minimum biomass required to get 80% of MMSY).

Location-specific recovery timeframes (Rt_i_) under a seascape moratorium scenario (i.e., minimum time for recovery to B^PGMY,low^, B_MMSY_ and B_PGMY,high_ based solely on fishing) were calculated by combining the posterior community growth rates with the location-specific standing stock biomass, unfished biomass and biomass target (i.e., B_PGMY,low_, B_MMSY_ or B_PGMY,high_) estimates (75). Following the logistic growth curve of biomass through time and assuming zero exports, we calculated the recovery time (Rt) as:

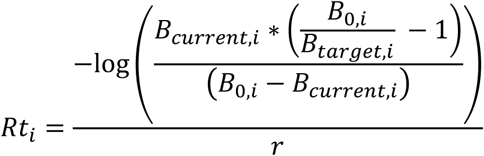

where *r* is the posterior biomass community growth rate, and B_current,i_, B_0,i_ and B_target,i_ are the posterior location-specific standing reef fish assemblage biomass, unfished biomass or target to be achieved (i.e., B_MMSY_ or B_PGMY_), respectively.

Note that recovery targets and timeframes could yield negative results for B_PGMY,low_ if the estimated biomass for a site or jurisdiction was below B_MMSY_ but above B_PGMY,low_.

We also estimated site-specific recovery times to B_MMSY_ using density-dependent harvesting scenarios (76), adding to the recovery time equation the harvest proportionality *h* (effort). We used a sequence of 10 harvesting values that varied from 0 (no fishing) to the sustainable exploitation rate that yields the multispecies surplus production (∼0.097), representing the minimum and maximum recovery times when harvesting for recovery (i.e., below the estimated surplus production).

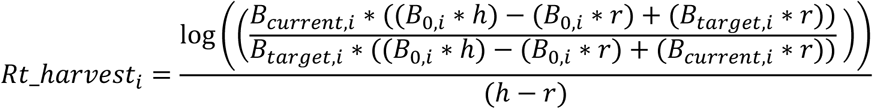

## Data and code availability

Data and code used for this paper is available for review and will be available from Github.com (JZamborain-Mason/ZamborainMasonetal_YieldGains).

## Acknowledgements

The authors would like to thank Ivor Williams, Rick Stuart-Smith, Graham Edgar, Shaun Wilson, Nicholas Polunin, Michel Kulbicki, Tim McClanahan, Pascale Chabanet, Eran Brokovich, Marah Hardt, Juan Cruz, Laurent Vigliola, Mark Tupper, and Stuart Sandin for providing data. We also thank all the data collectors including the NOAA Coral Reef Ecosystems Division (funding by NOAA Coral Reef Conservation Program).

## Author contributions

JZM conceived the study, developed and implemented the analyses, and wrote the manuscript with support from SRC, JEC, and MAM. All other authors contributed data and/or made substantive contributions to the text.

## Classification

Biological Sciences (Agricultural Sciences)/Social Sciences (Sustainability Science)

## Supporting information

**Figure S1.**
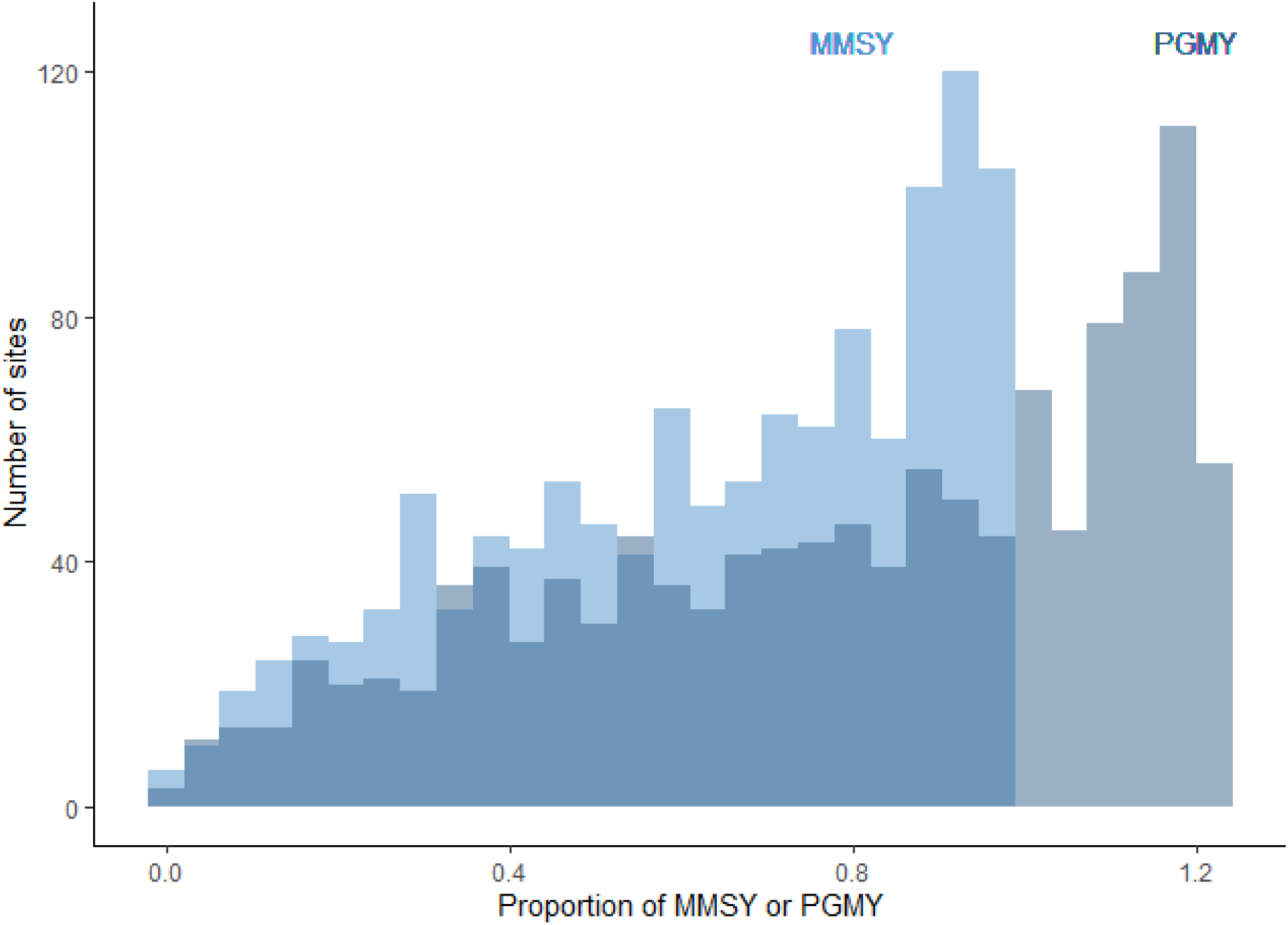
Frequency of site-specific sustainable yields relative to MMSY or PGMY. Estimates are best-fit median estimates (i.e., median site-specific surplus divided by median site-specific MMSY or PGMY; similar to the food provision index in ref (28)).

**Figure S2.**
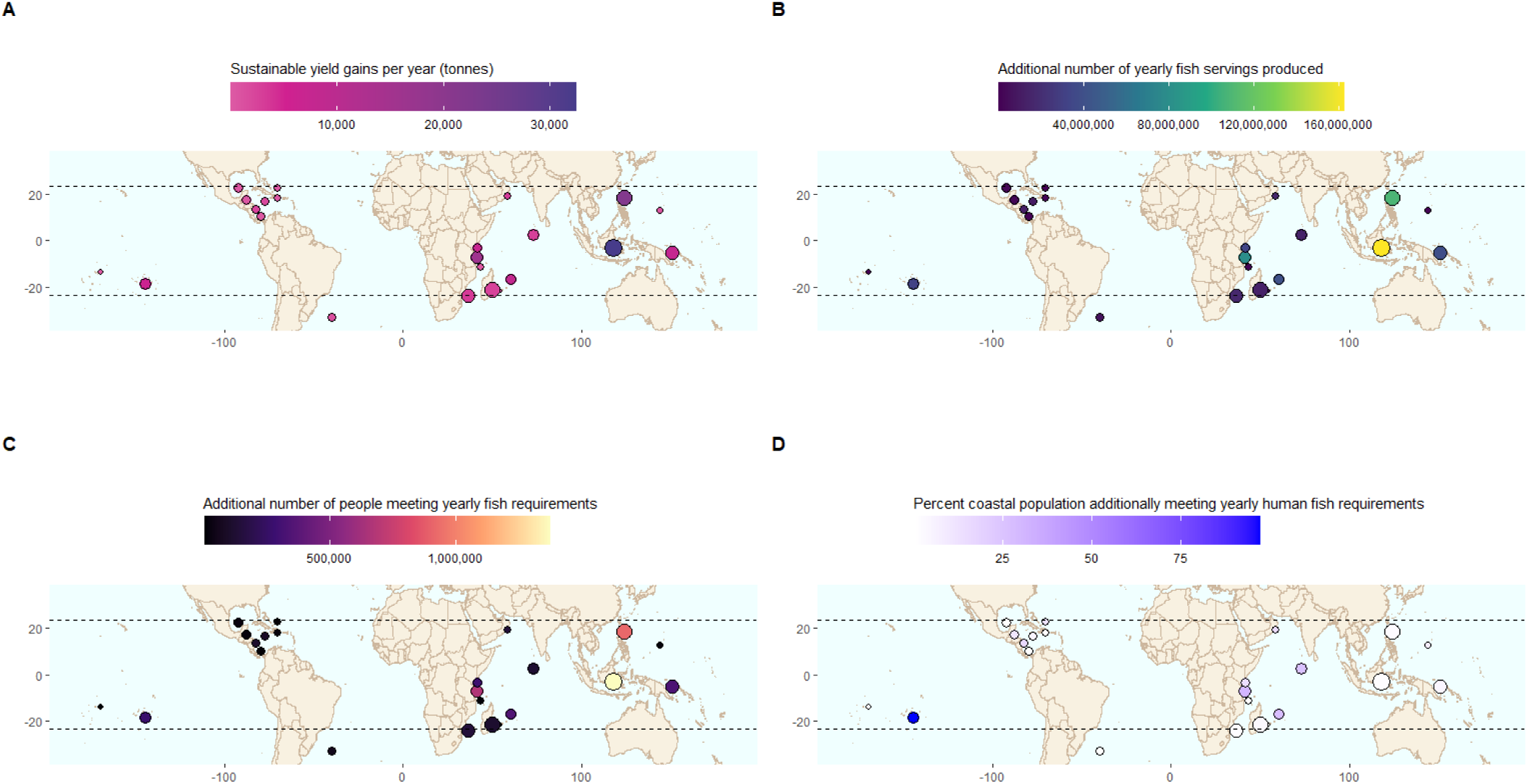
Potential jurisdiction yearly gains based on their estimated reef area open to fishing. **(A)** Sustainable yield gains. (**B**) additional number of 100 g fish servings produced. (**C**) additional number of people meeting yearly fish intake recommendations. (**D**) additional number of people meeting yearly fish intake recommendations as a percentage of the coastal population, defined as population living within 30km of the coast (17, 72, 73). Point size in **A-D** is scaled according to the estimated reef area open to fishing

**Figure S3.**
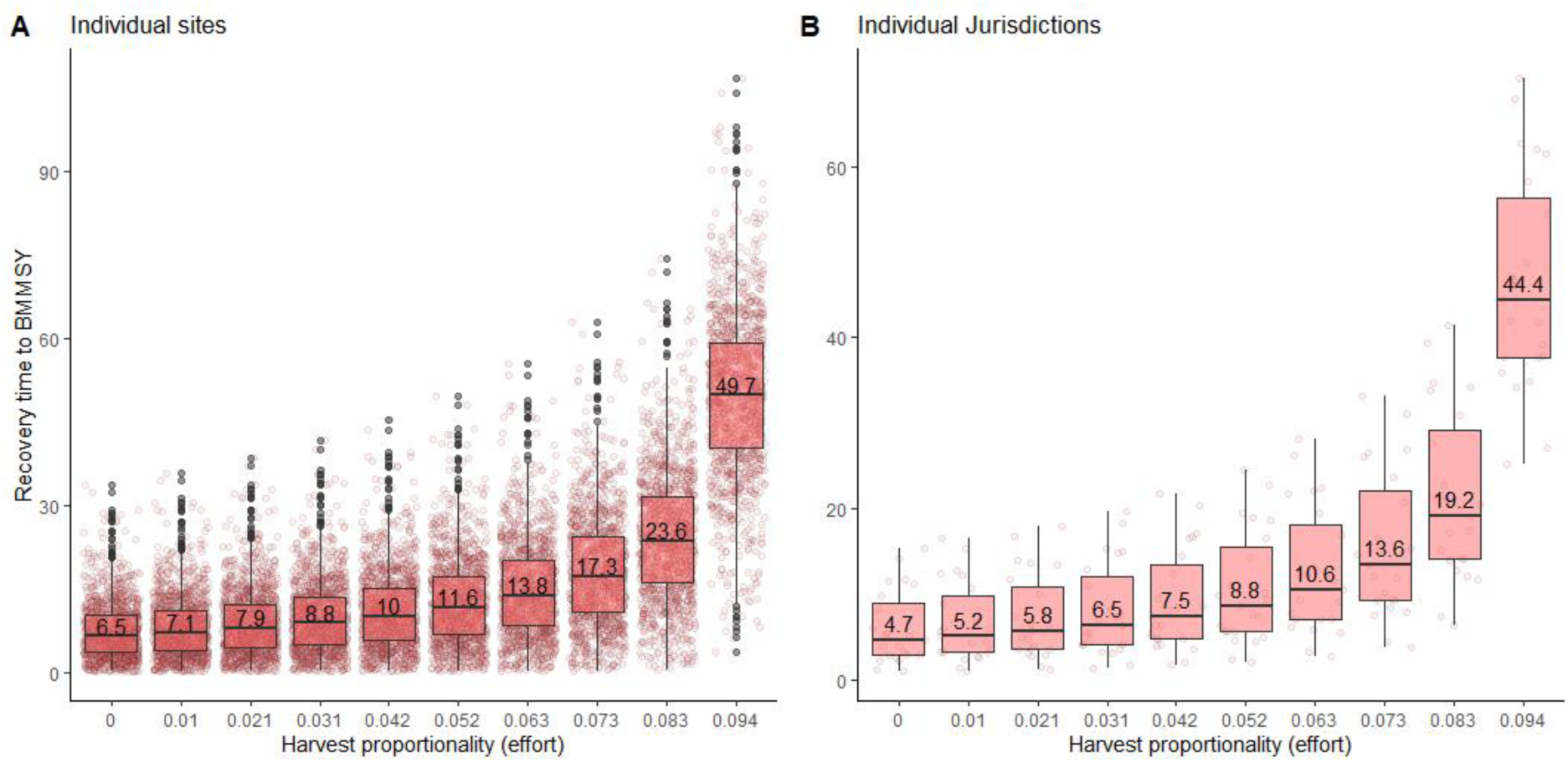
Estimated recovery times (years) to B_MMSY_ under different seascape density-dependent harvesting scenarios. **(A)** For individual reef sites (each point); and **(B)** for individual jurisdictions (each point). Numbers are median recovery times across sites **(A)** or jurisdictions **(B)**.

**Figure S4.**
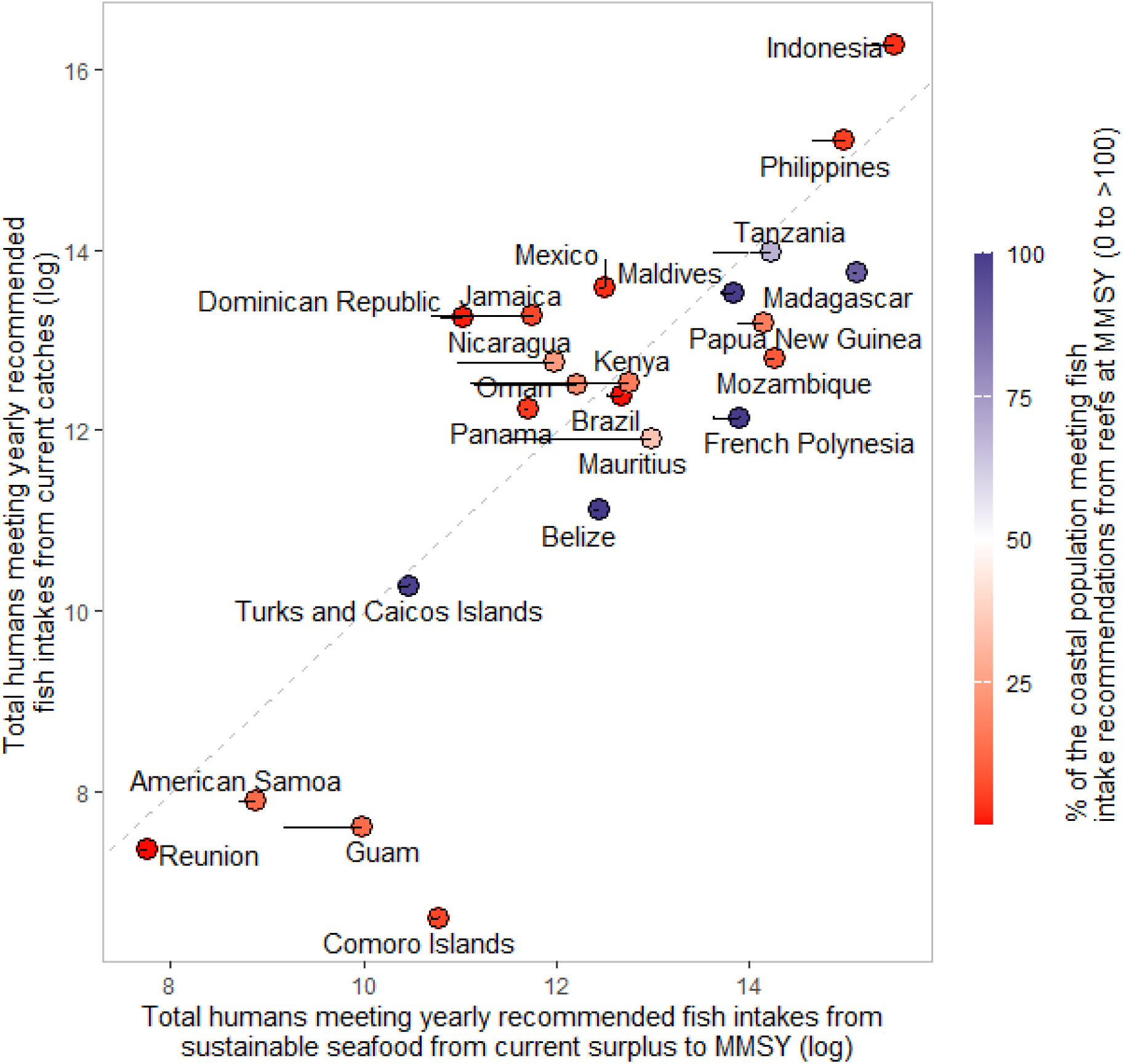
Reconstructed reef fish catches vs sustainable production in terms of total humans meeting yearly recommended fish intakes. Each point is a jurisdiction. The horizontal lines connecting points represent the total humans that could potentially meet yearly fish intake recommendations from the surplus (based on their standing biomass) to their MMSY (the actual point). The dashed line is the unity, representing the limit for jurisdictions that are catching above their estimated maximum production (i.e., those whose points are above the unity line).

**Table S1.**
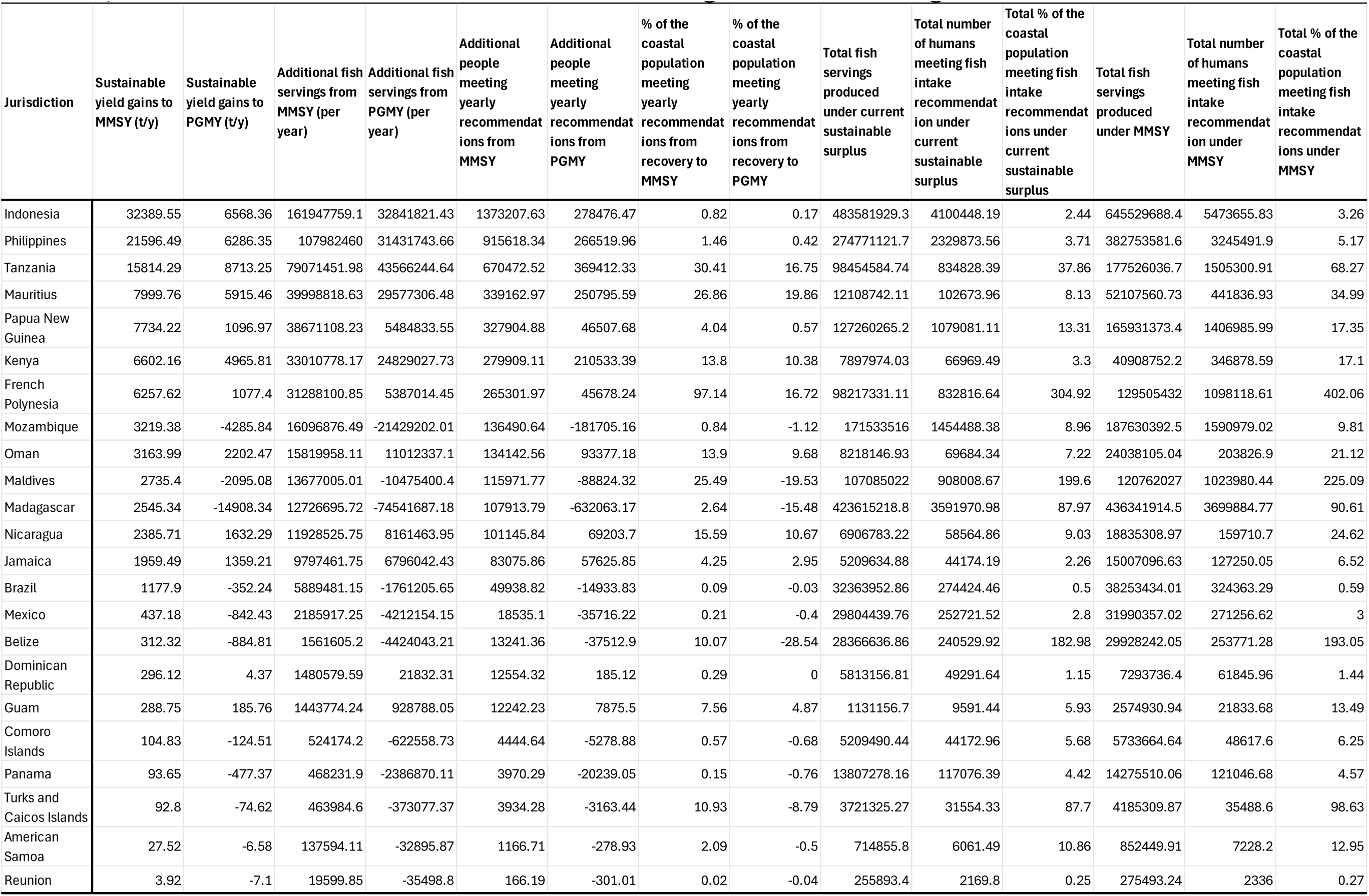
Jurisdiction-level estimated benefits from recovering reef fish assemblages. Values are based on median values.

**Table S2.** Reef families included in our analyses.

| **Fish family** | **Common family name** |
| --- | --- |
| Acanthuridae | Surgeonfishes |
| Balistidae | Triggerfishes |
| Caesionidae | Fusiliers |
| Carangidae | Jacks |
| Chaetodontidae | Butterflyfishes |
| Cirrhitidae | Hawkfishes |
| Diodontidae | Porcupinefishes |
| Ephippidae | Batfishes |
| Haemulidae | Sweetlips |
| Kyphosidae | Drummers |
| Labridae | Wrasses and Parrotfishes |
| Lethrinidae | Emperors |
| Lutjanidae | Snappers |
| Monacanthidae | Filefishes |
| Mullidae | Goatfishes |
| Nemipteridae | Coral Breams |
| Pinguipedidae | Sandperches |
| Pomacanthidae | Angelfishes |
| Serranidae | Groupers |
| Siganidae | Rabbitfishes |
| Sparidae | Porgies |
| Sphyraenidae | Barracudas |
| Synodontidae | Lizardfishes |
| Tetraodontidae | Pufferfishes |
| Zanclidae | Moorish Idol |

